## Supplementary Information for "The Therapeutic Nanobody Profiler: characterising and predicting nanobody developability to improve therapeutic design"

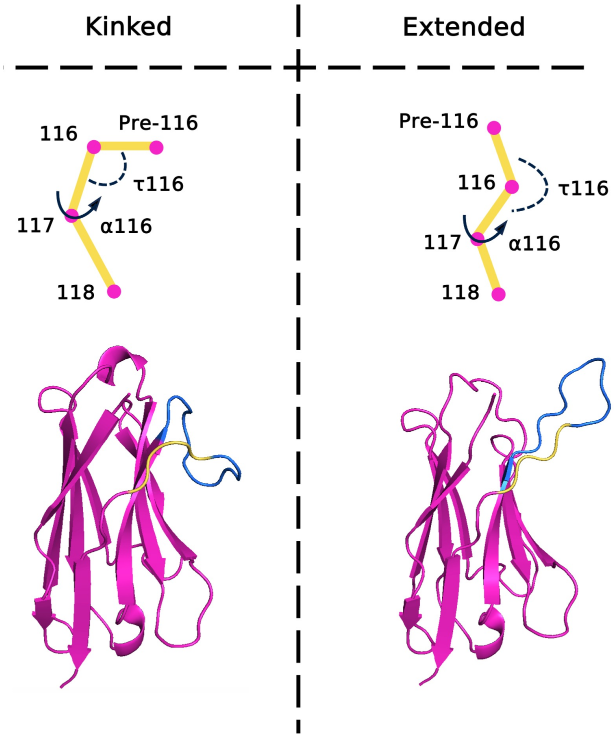


**Figure S1.** CDR3 conformations can be classified as either ‘kinked’ or ‘extended’, following methods in Weitzner *et al.* (2015) and Bahrami-Dizicheh *et al.* (2023) [45, 46]. This classification is based on the $\tau$ pseudo-dihedral and $\alpha$ angles between backbone C$\alpha$ atoms in the CDR3 at positions 116, 117 and 118 (by IMGT numbering). Kinked loops have $\alpha$ and $\tau$ angles in the range: $0° < \alpha 116 < 120°$, $85° < \tau 116 < 130°$. Extended loops have $\alpha$ and $\tau$ angles in the range: $-100° > \alpha 116$, $100° < \tau 116 < 145°$. Figure is adapted from Bahrami-Dizicheh *et al.* [46].

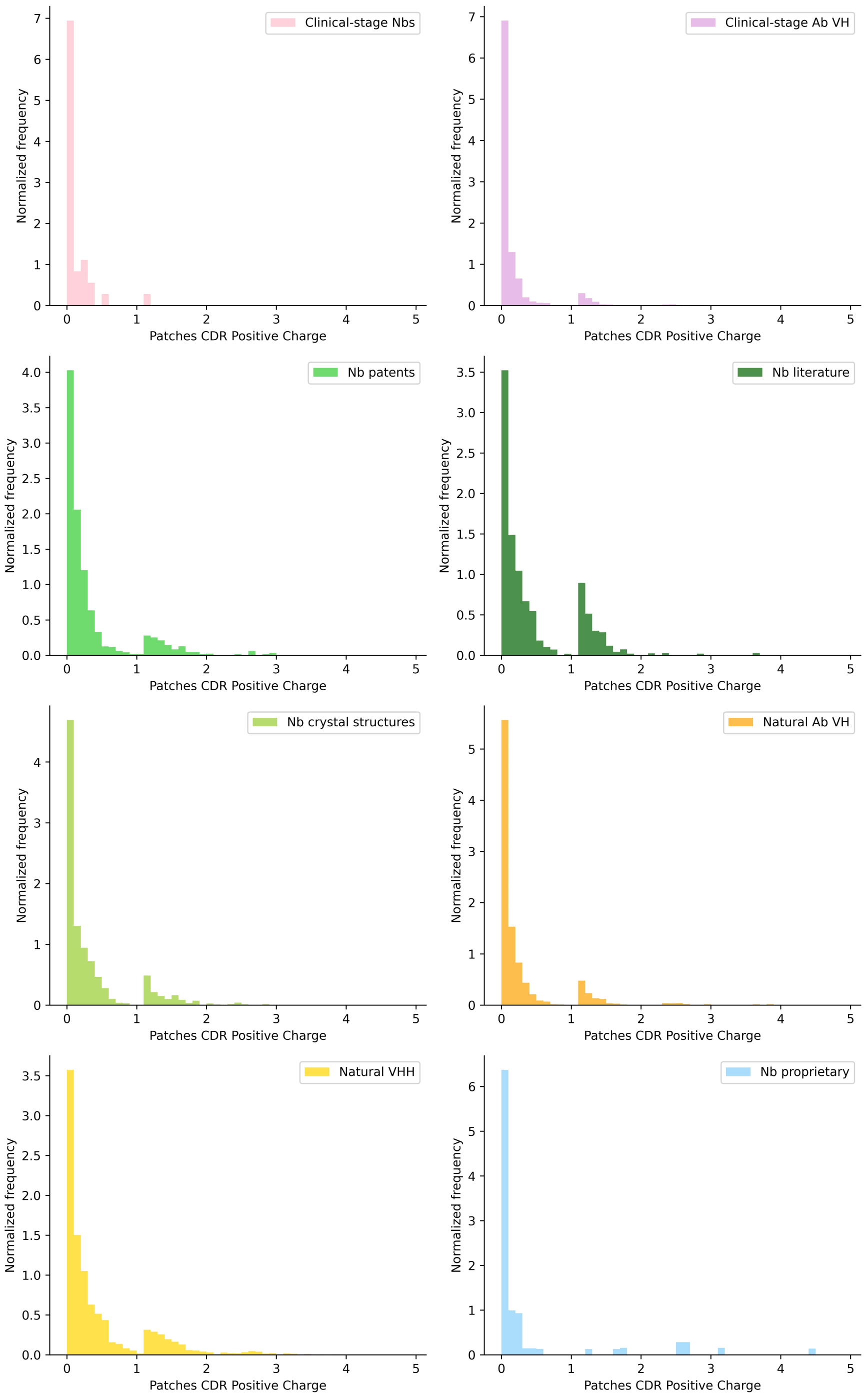


**Figure S2.** Distributions of scores for patches of positive charge across all our datasets, determined by sampling with replacement all datasets (aside from the 36 clinical-stage nanobodies) with sample size of 36 and 300 samples.

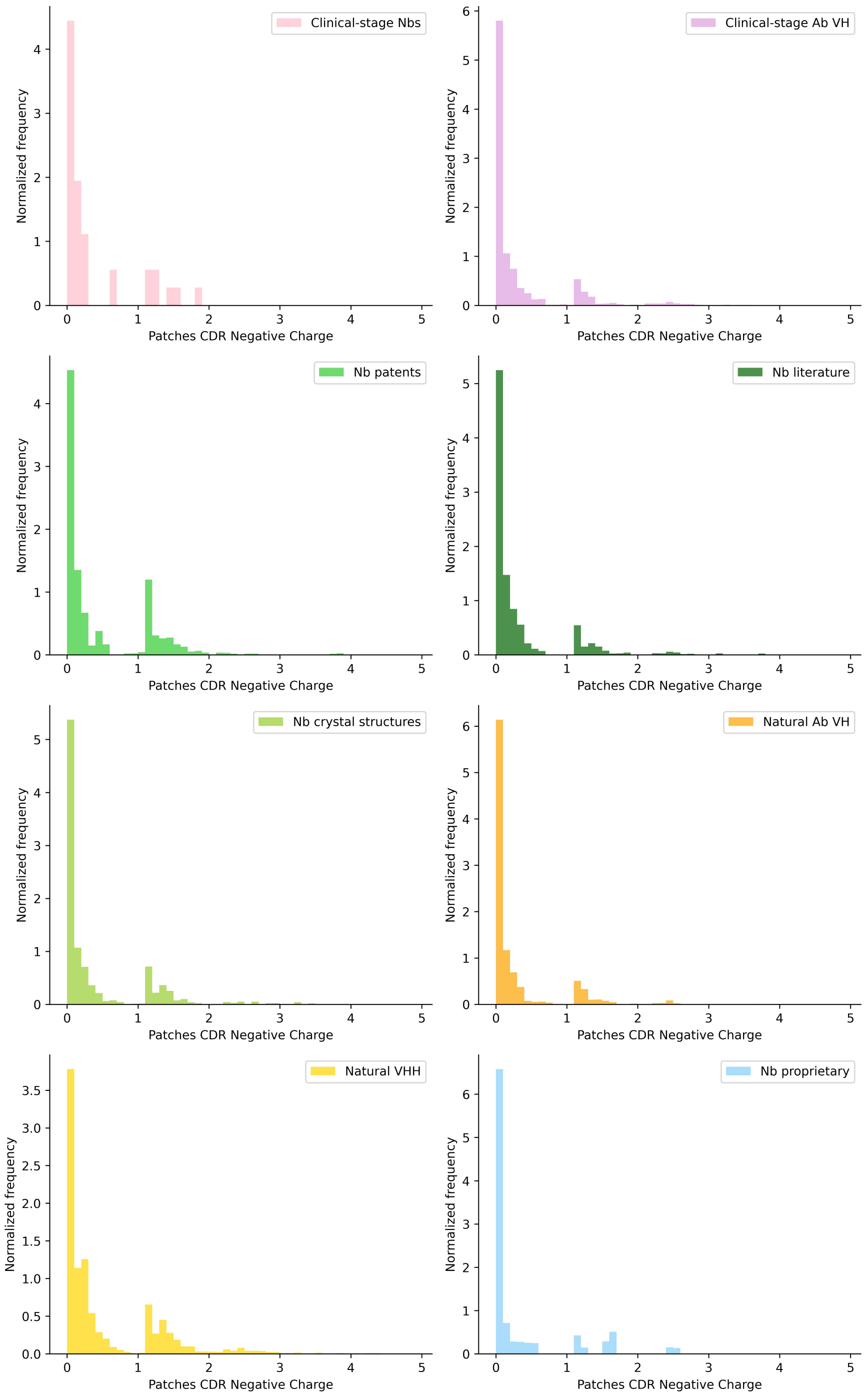


**Figure S3.** Distributions of scores for patches of negative charge across all our datasets, determined by sampling with replacement all datasets (aside from the 36 clinical-stage nanobodies) with sample size of 36 and 300 samples.

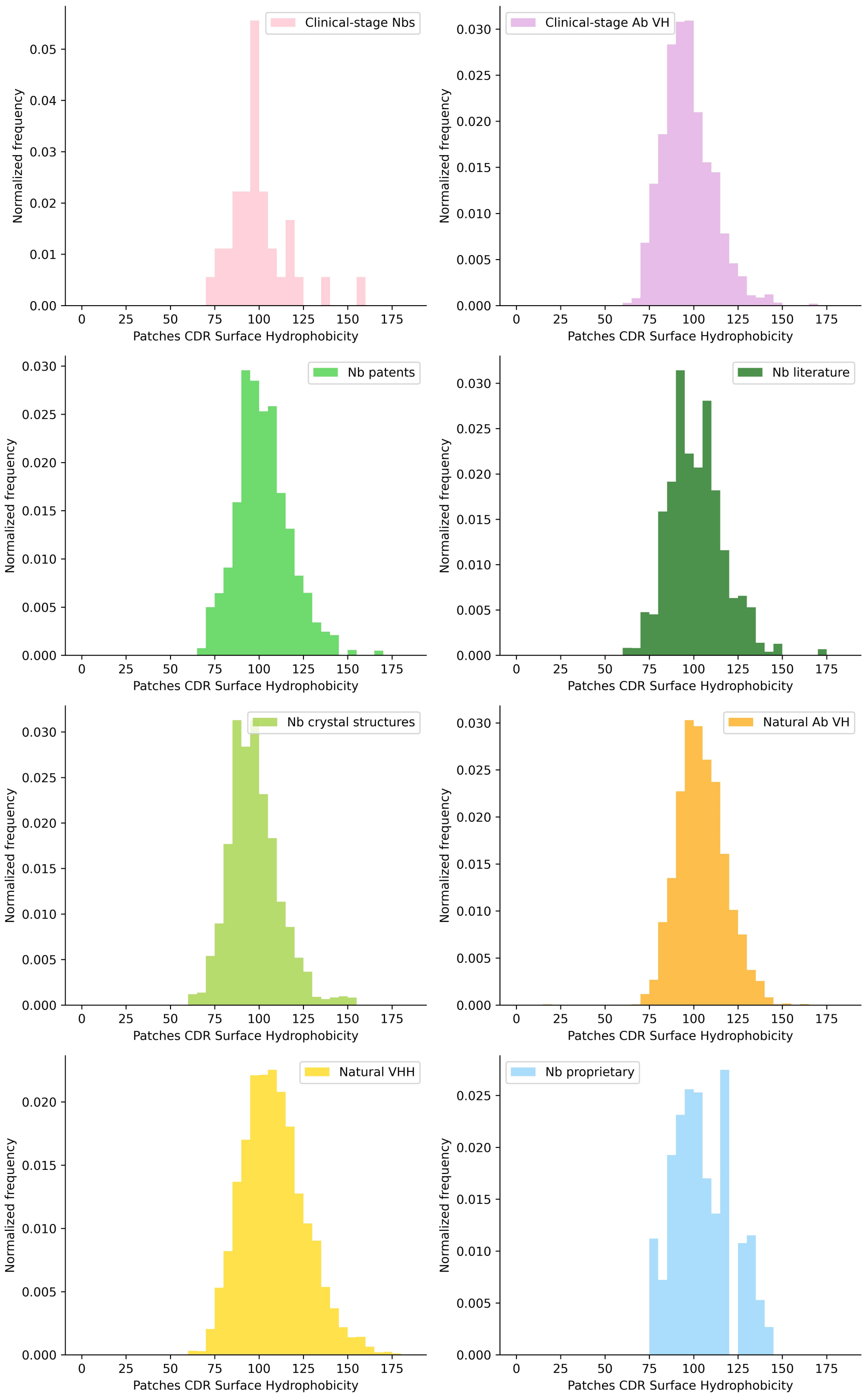


**Figure S4.** Distributions of scores for patches of surface hydrophobicity across all our datasets, determined by sampling with replacement all datasets (aside from the 36 clinical-stage nanobodies) with sample size of 36 and 300 samples.

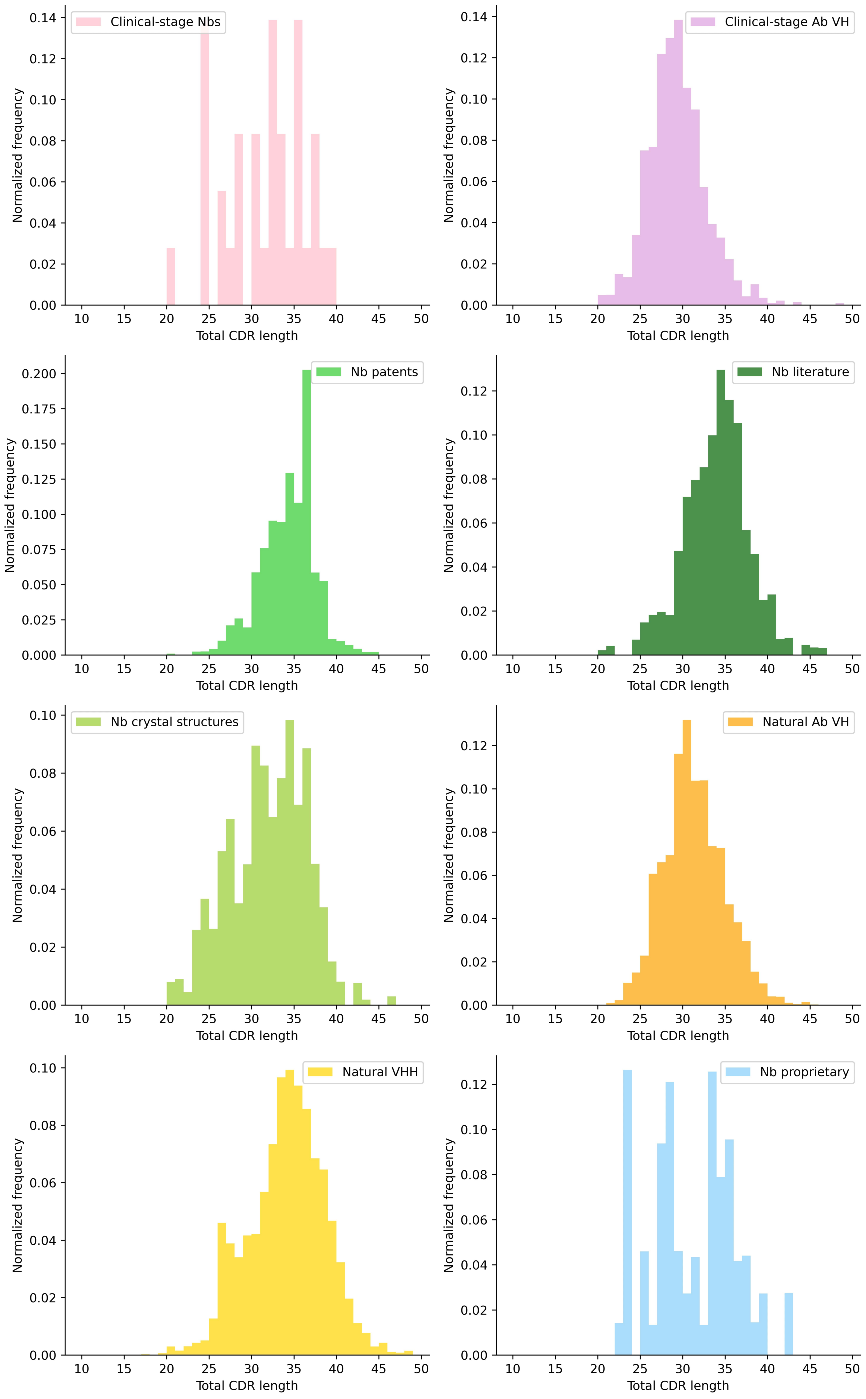


**Figure S5.** Distributions of total CDR length across all our datasets, determined by sampling with replacement all datasets (aside from the 36 clinical-stage nanobodies) with sample size of 36 and 300 samples.

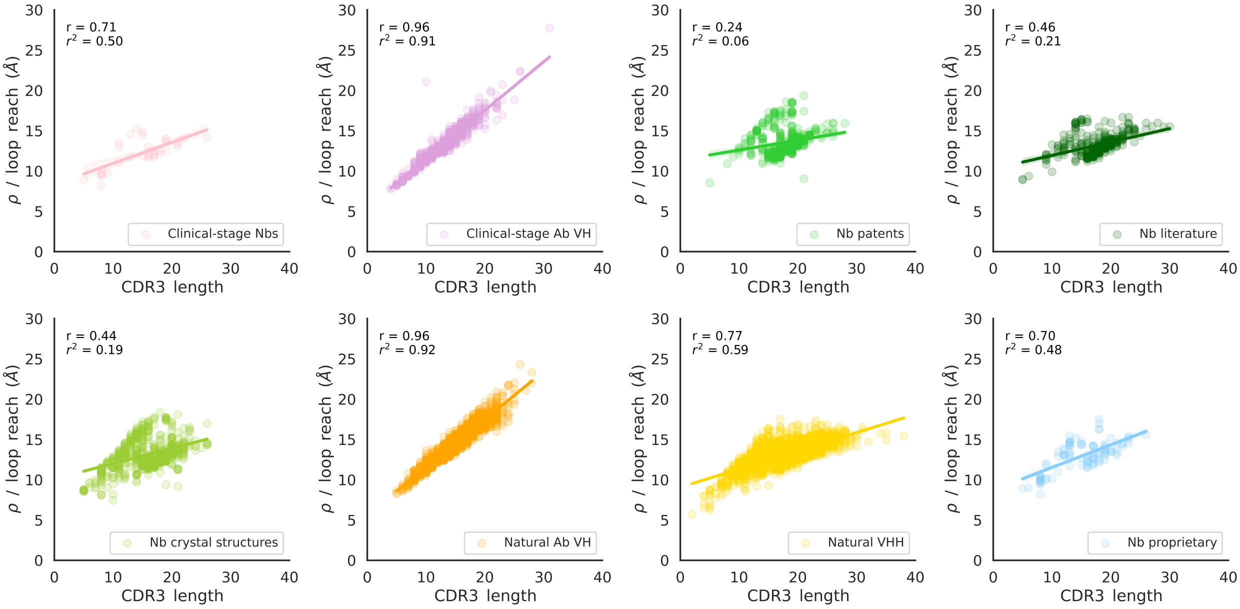


**Figure S6.** Correlations between reach and length of the CDR3 loop for the different datasets analysed in this study show divergent trajectories for nanobody data, where an increase in length does not translate to greater reach. This relationship can be quantified by loop compactness.

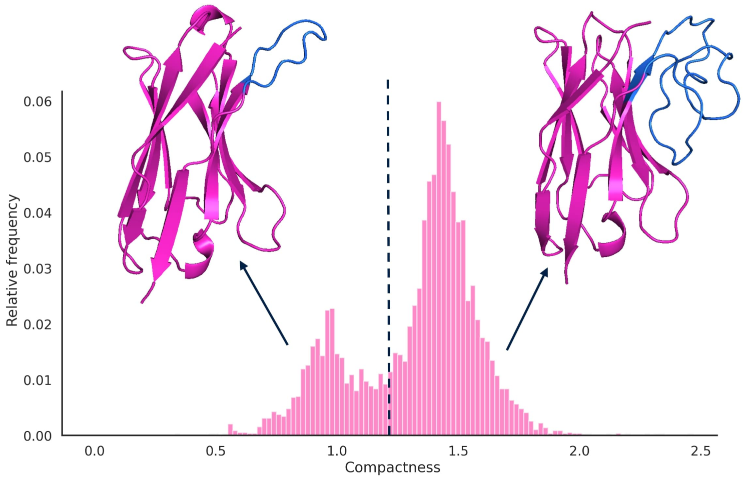


**Figure S7.** Loop compactness provides a continuous descriptor for CDR3 conformation. The two peaks in the bimodal distribution of compactness for all of our nanobody datasets agree with previous classifications of loop conformations as either extended (lower compactness) or kinked (higher compactness).

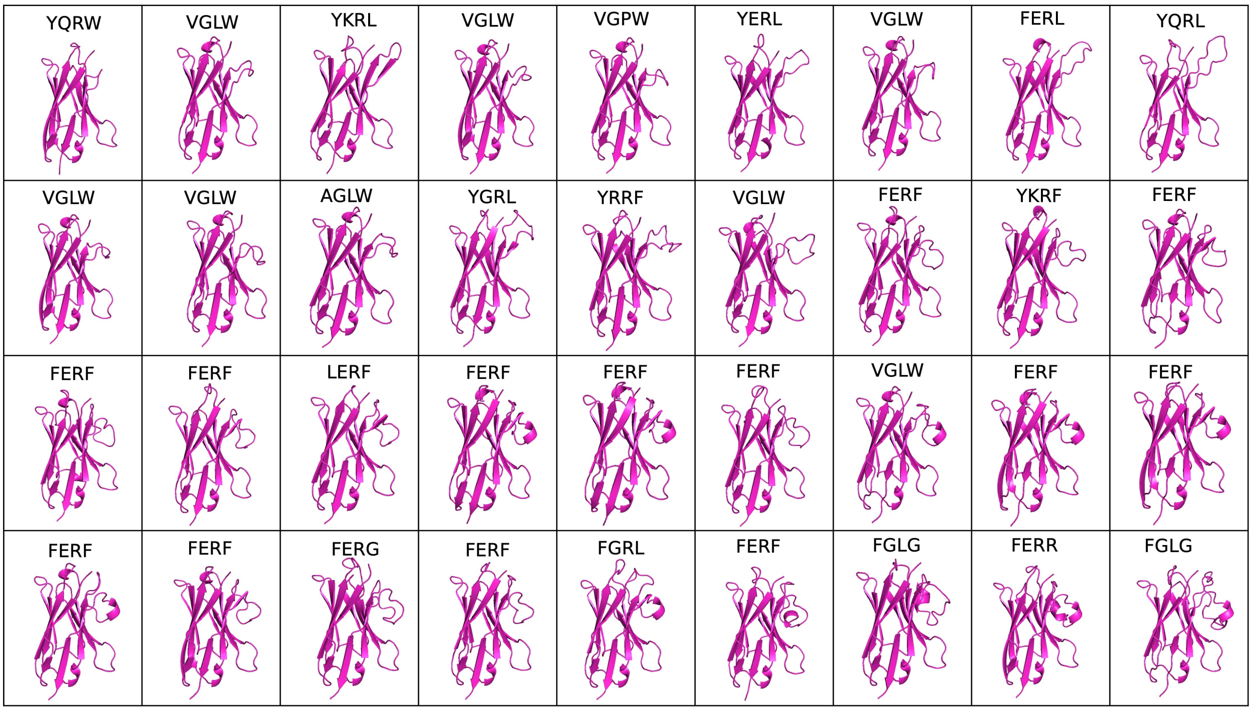


**Figure S8.** Predicted structures for 36 clinical-stage nanobody sequences in order of increasing compactness (top to bottom, left to right) alongside nanobody tetrad motifs (amino acid identities at IMGT positions 42, 49, 50, 52). As compactness increases, so does CDR3 length and conformational complexity.

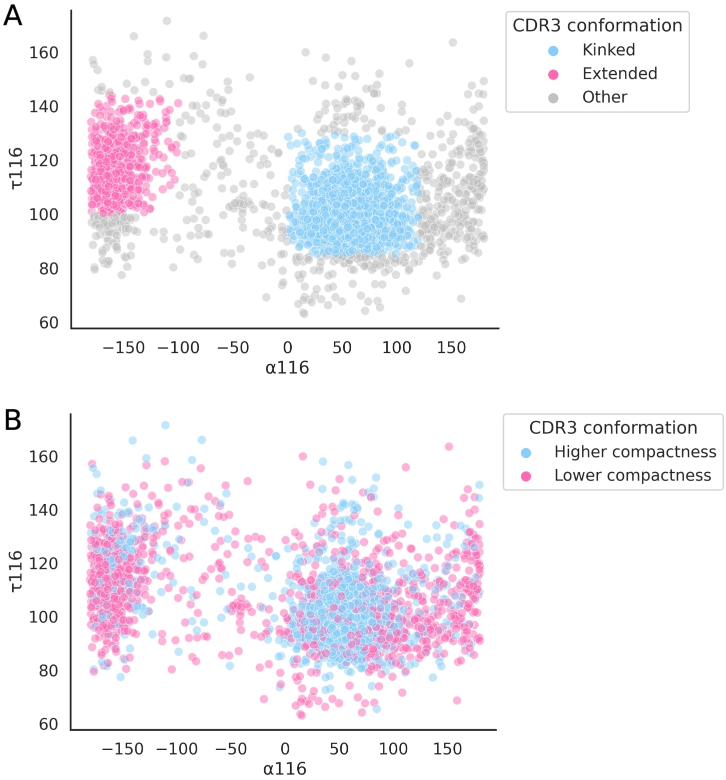


**Figure S9.** CDR3 conformations can be classified as kinked or extended according to the alpha and tau angles between backbone atoms around position 116 (IMGT numbering) in the CDR3 loop [45, 46]. (**A**) Classification of the CDR3 conformation for all nanobodies from our datasets combined shows that many do not fall into either category using this method. (**B**) Alternatively the CDR3 conformation may be described using compactness. Using the same method to cluster the nanobodies on CDR3 angles, divided into groups of higher and lower compactness, shows the limitations of the classification method given the large conformational diversity.

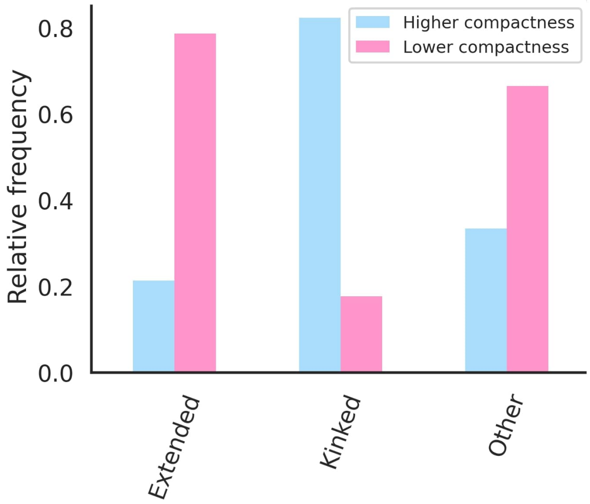


**Figure S10.** To assess the agreement between compactness and the kinked-extended classifications, a threshold was drawn between the two peaks of the bimodal distribution (**Figure S7**) at 1.25 to separate the data into loops of lower versus higher compactness. Loops with lower compactness closely align with those classified as extended, while those with higher compactness correspond to kinked loops, showing consistency between the compactness measure and the classification method in describing CDR3 conformation.

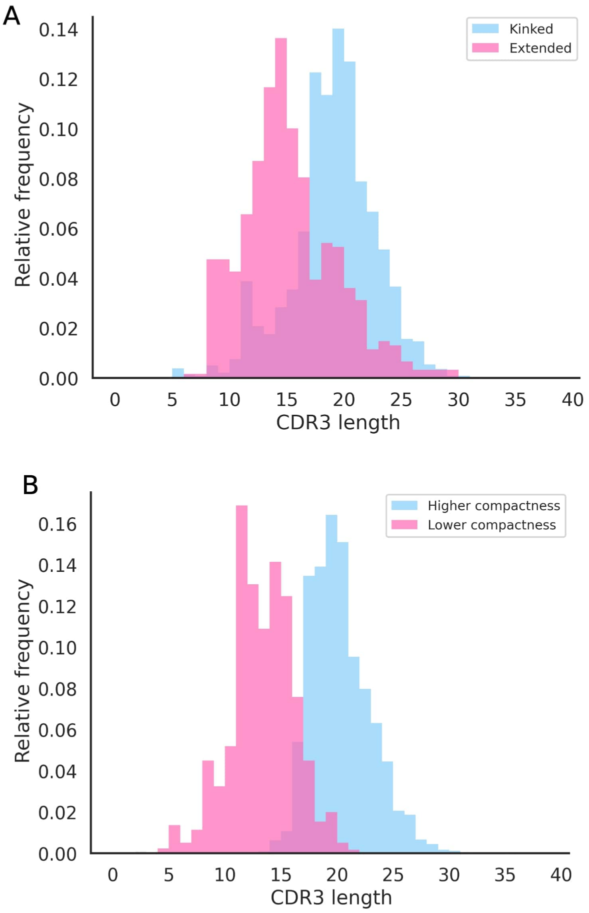


**Figure S11.** The bimodal distribution for compactness can be used to separate nanobodies into two structural subtypes. We find that these demonstrate distinguishable profiles of secondary features associated with the CDR3 loop structure which are consistent across descriptors for CDR3 conformation. When comparing CDR3 loop length, (**A**) CDR3 loops with a kinked conformation tend to be longer in terms of number of residues than extended conformations. (**B**) CDR3 loops with a higher compactness score tend to be longer than loops of lower compactness.

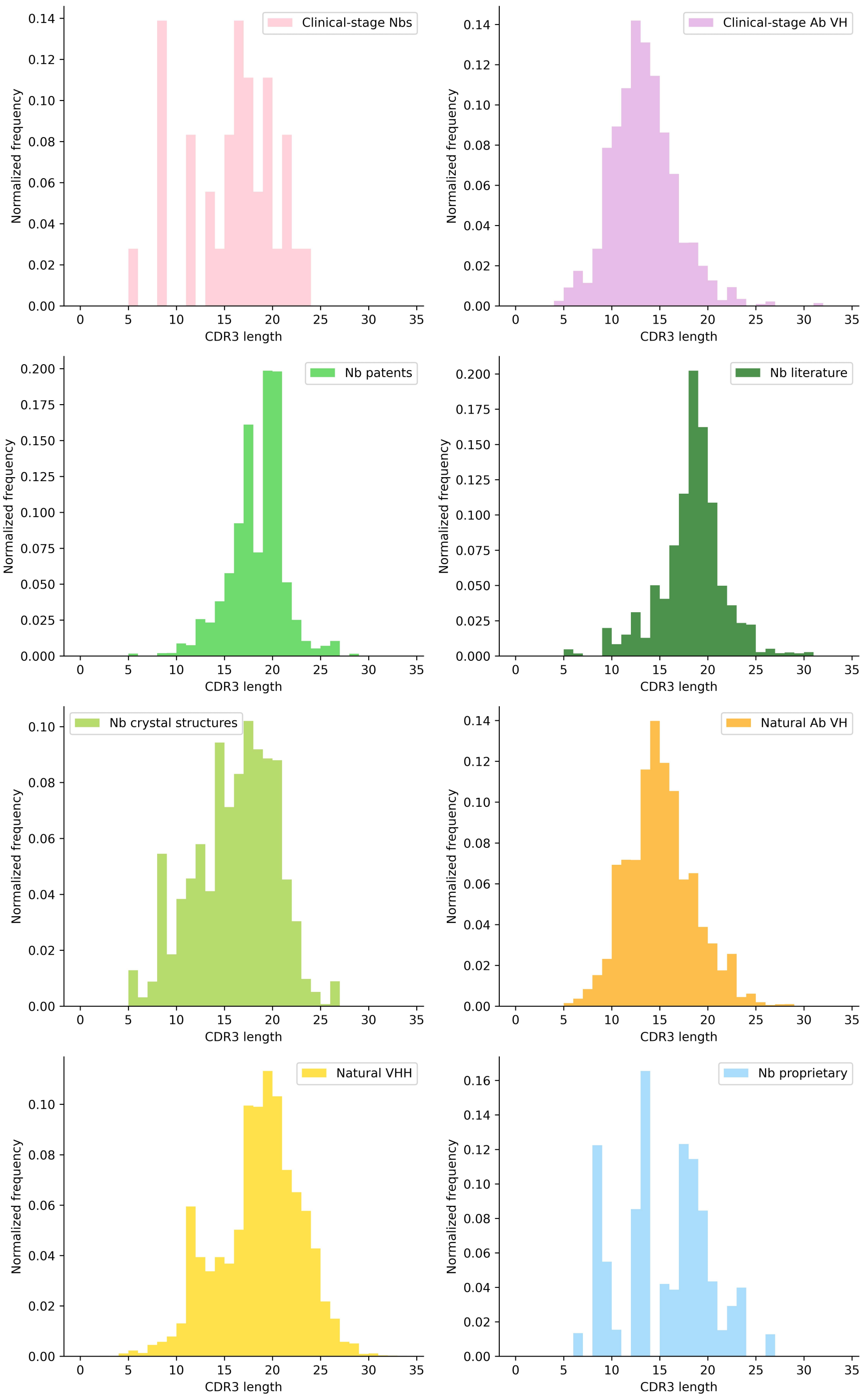


**Figure S12.** Distributions of CDR3 length across all our datasets, determined by sampling with replacement all datasets (aside from the 36 clinical-stage nanobodies) with sample size of 36 and 300 samples.

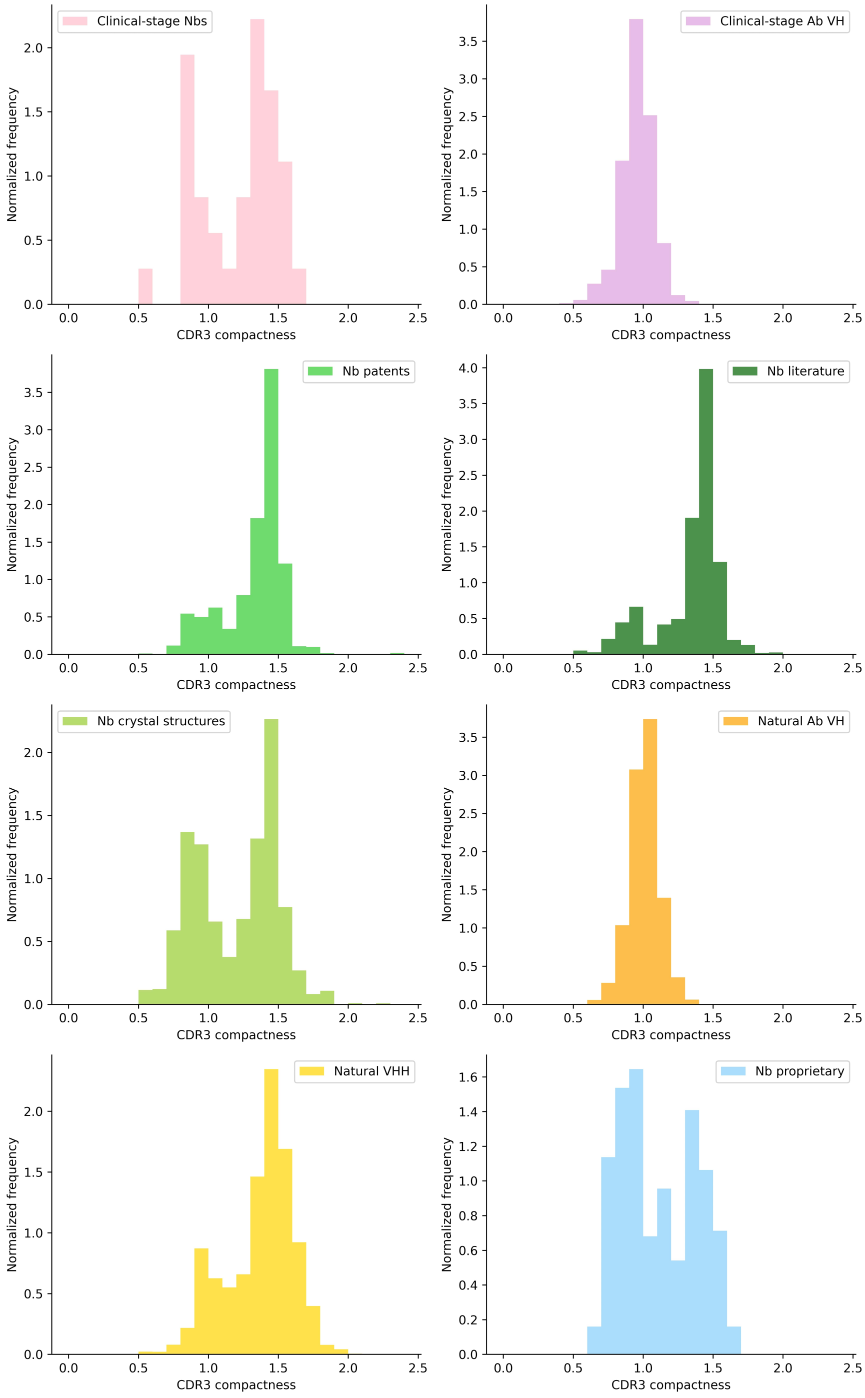


**Figure S13.** Distributions of CDR3 compactness scores across all our datasets, determined by sampling with replacement all datasets (aside from the 36 clinical-stage nanobodies) with sample size of 36 and 300 samples.

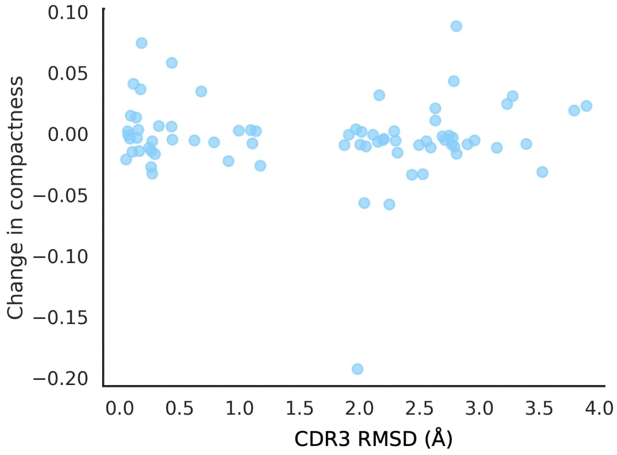


**Figure S14.** Change in compactness is minimal between bound and unbound pairs of nanobody CDR3 loop structures, and does not show a positive correlation with the RMSD over the bound and unbound CDR3 loops.

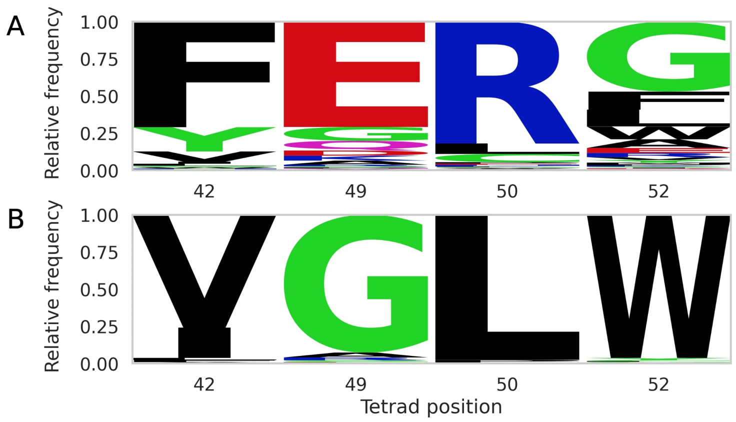


**Figure S15.** Proportions of amino acids across nanobody tetrad (positions 42, 49, 50 and 52 by IMGT numbering) in our (**A**) nanobody versus (**B**) antibody VH datasets shows that FERG is the most common nanobody motif, whilst VGLW is the most common in antibodies.

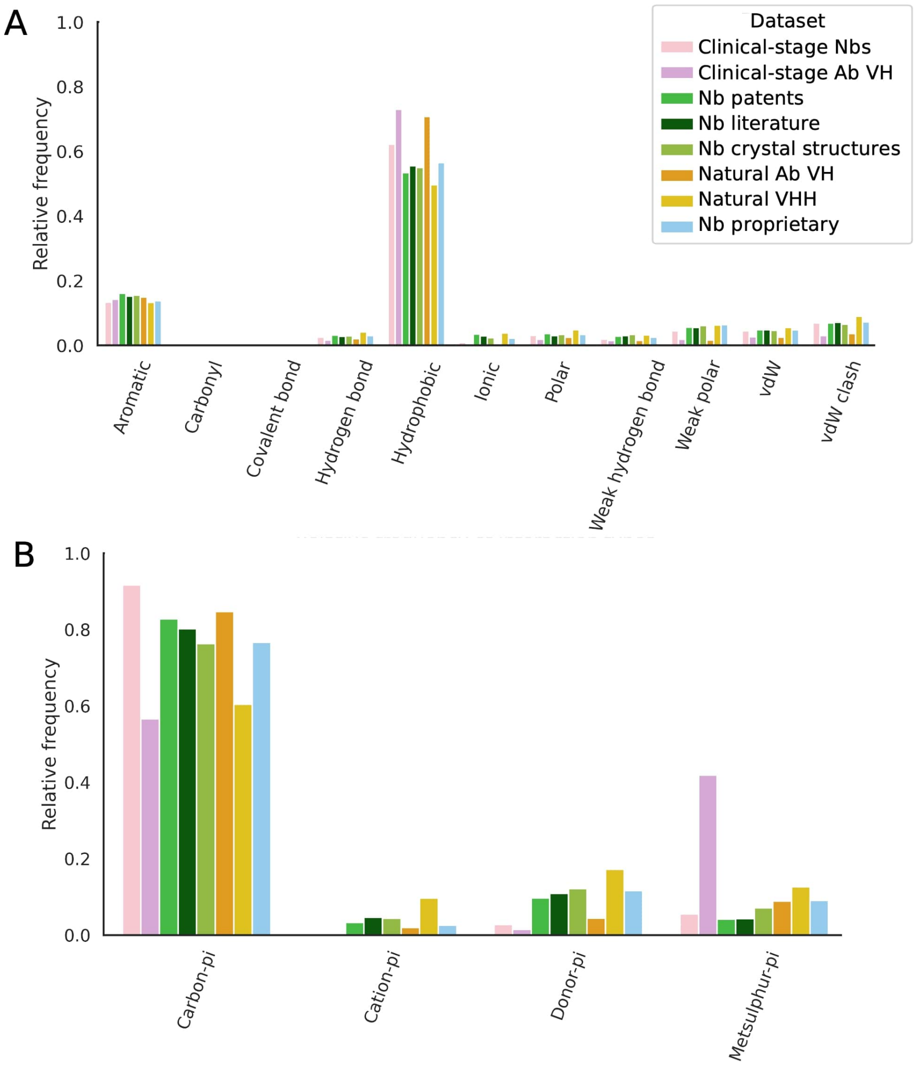


**Figure S16.** Of pairs of residues in the CDR3 loop and the tetrad positions across all nanobody datasets that were found to interact, for (**A**) atom-atom interactions, hydrophobic, and aromatic interactions were most common. For (**B**) atom-plane interactions, carbon-$\pi$ interactions were most common.

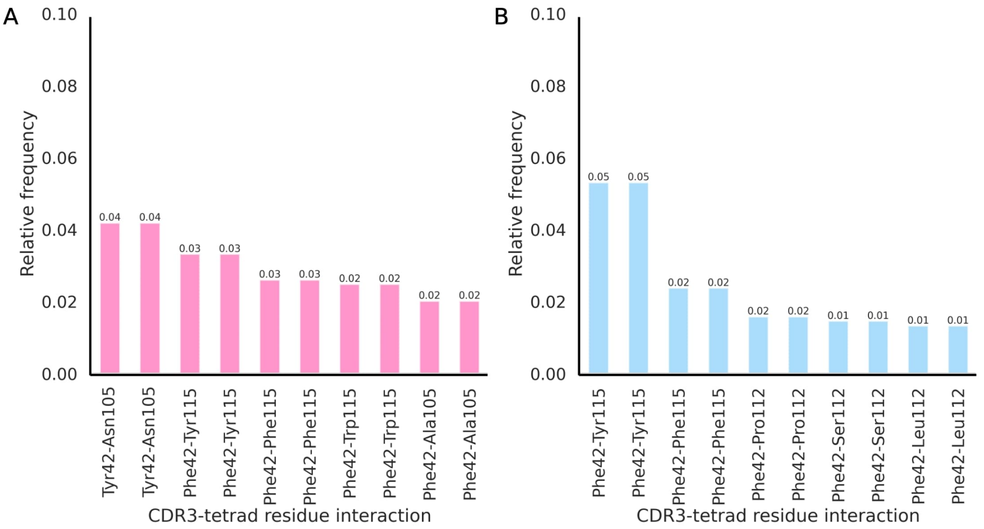


**Figure S17.** Top 10 most common interacting CDR3-tetrad residue pairs in decreasing order of relative frequency across all nanobody datasets for (**A**) those with CDR3 loops of lower compactness (that tend to extend out from the framework) and (**B**) loops of higher compactness (that tend to fold over the FR2 region).

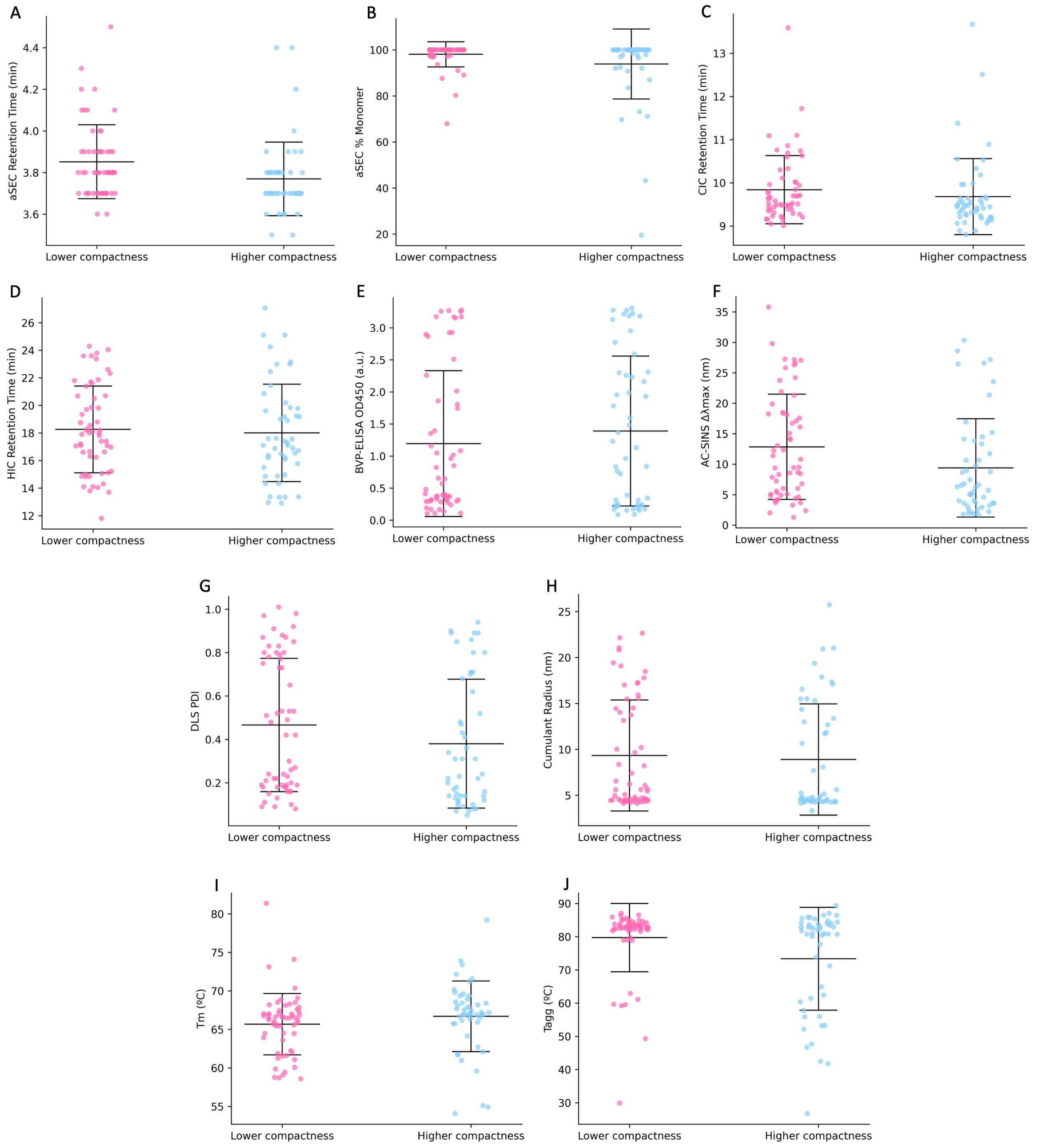


**Figure S18.** Results of experimental assays split into nanobodies with CDR3 loops of higher and lower compactness at a compactness score threshold of 1.25. (**A**) aSEC retention time (**B**) aSEC % monomer (**C**) CIC retention time (**D**) HIC retention time (**E**) BVP-ELISA OD450 (**F**) AC-SINS $\Delta\lambda$max (**G**) DLS PDI (**H**) Cumulant radius (**I**) Tm (melting temperature) (**J**) Tagg (aggregation temperature)

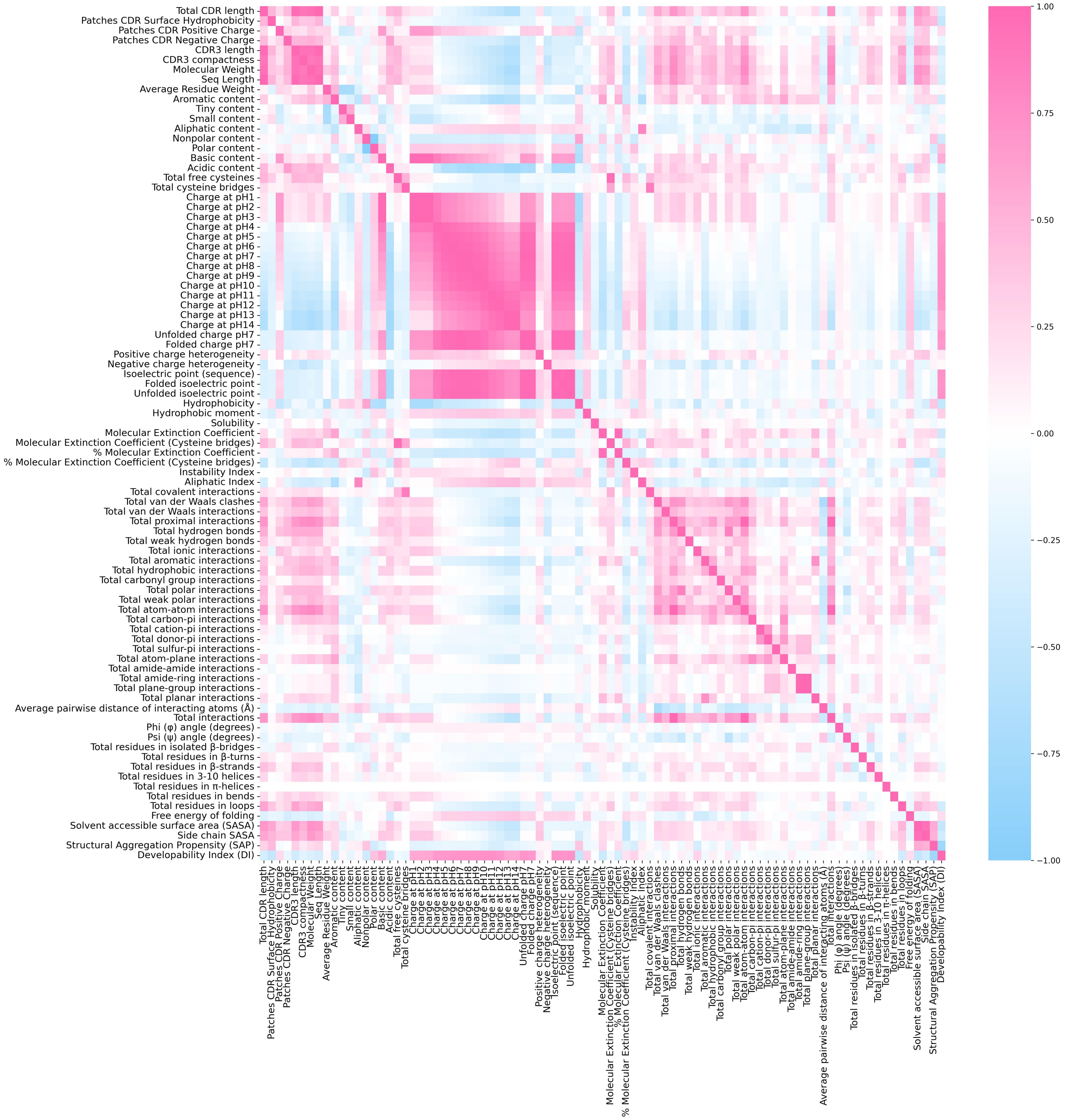


**Figure S19.** Heatmap of Spearman's rank correlations between all computational metrics calculated for the combined nanobody datasets only.

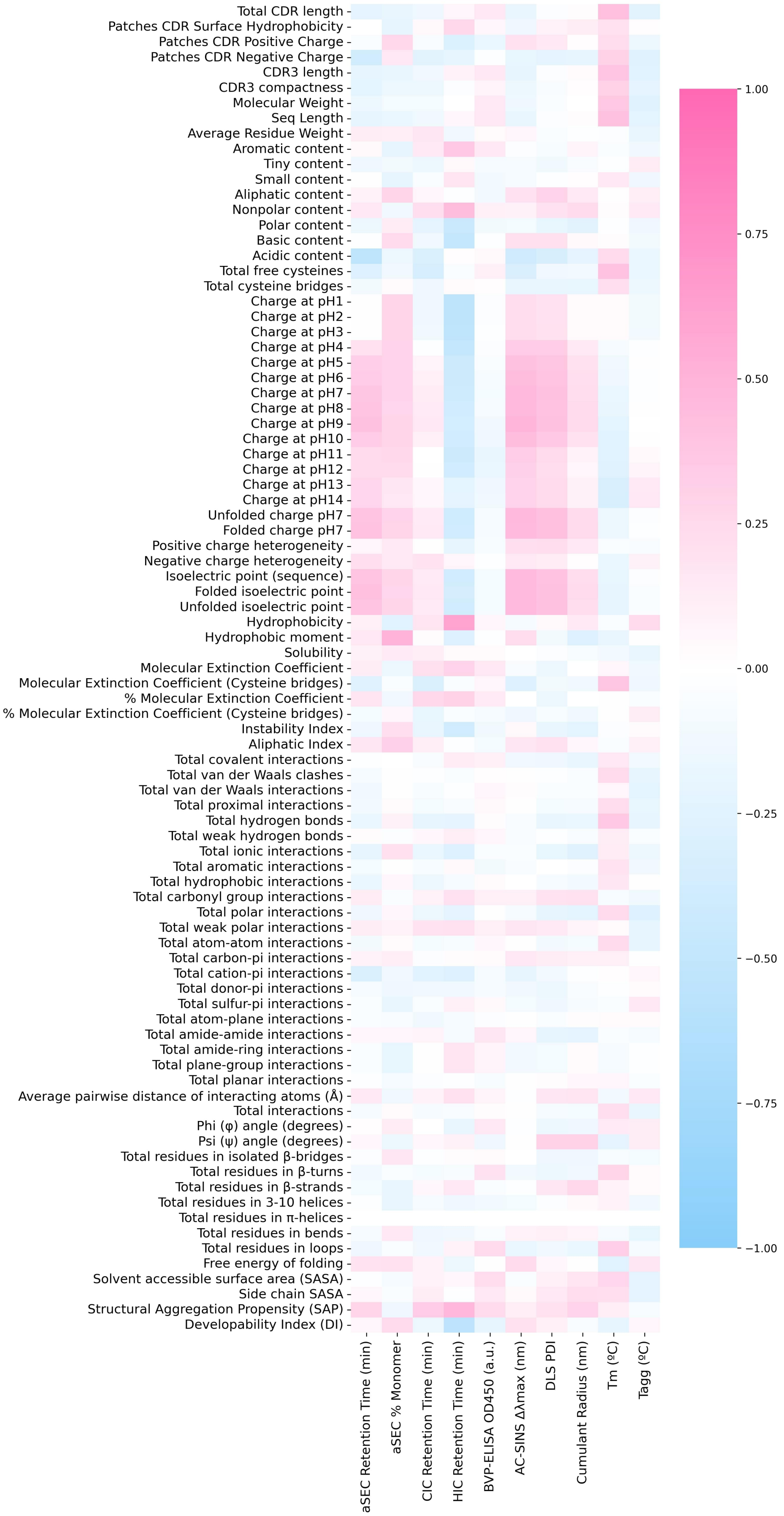


**Figure S20.** Heatmap of Spearman's rank correlations between all computational metrics calculated for the clinical-stage and proprietary nanobody datasets only against their *in vitro* assay results.

| | Unbound PDB | Bound PDB | | Unbound PDB | Bound PDB | | Unbound PDB | Bound PDB |
|---|---|---|---|---|---|---|---|---|
| 1 | 8f8v | 8f8w | 26 | 5tp3 | 5toj | 51 | 7np9 | 7p2d |
| 2 | 8f8v | 8f8x | 27 | 4krn | 4kro | 52 | 6dbg | 6dbf |
| 3 | 8gz6 | 8gz5 | 28 | 5h8d | 5h8o | 53 | 6dba | 6dbg |
| 4 | 7sah | 6lr7 | 29 | 6y0e | 6xzf | 54 | 6dba | 6dbf |
| 5 | 7vpy | 7vq0 | 30 | 3r0m | 7ri2 | 55 | 6apq | 4w2o |
| 6 | 4b50 | 7aej | 31 | 3r0m | 3rjq | 56 | 5wb1 | 4xt1 |
| 7 | 6u53 | 6u54 | 32 | 5sv4 | 5sv3 | 57 | 5wb2 | 4xt1 |
| 8 | 4w2p | 4w2q | 33 | 4s11 | 4s10 | 58 | 5hdo | 5hgg |
| 9 | 5vm4 | 5vm6 | 34 | 4s11 | 6h1f | 59 | 5lz0 | 6ey0 |
| 10 | 7kgl | 7kgk | 35 | 4qgy | 4y7m | 60 | 6u50 | 6u51 |
| 11 | 7mfv | 7kgk | 36 | 4m3j | 4m3k | 61 | 6u50 | 6u52 |
| 12 | 7lvu | 7lvw | 37 | 5m2w | 5mwn | 62 | 7om5 | 7om4 |
| 13 | 5e7b | 5e7f | 38 | 5m2w | 5m30 | 63 | 7omt | 7omm |
| 14 | 6fys | 6fyw | 39 | 7kkj | 7kkl | 64 | 2xa3 | 7r73 |
| 15 | 6ksn | 7d5p | 40 | 6ck8 | 6fyt | 65 | 6cwk | 6cwg |
| 16 | 6ksn | 7d5q | 41 | 6z1z | 6z20 | 66 | 6u14 | 6u12 |
| 17 | 3qxu | 3qxt | 42 | 5vlv | 5vm0 | 67 | 6ru3 | 6ruv |
| 18 | 7v0v | 3k1k | 43 | 3qxw | 7rga | 68 | 6ru3 | 7noz |
| 19 | 7v0v | 3ogo | 44 | 3qxw | 7rg7 | 69 | 2xxc | 2xt1 |
| 20 | 1f2x | 1g6v | 45 | 3qxw | 3qxv | 70 | 2xxc | 2xv6 |
| 21 | 6t2j | 6sc5 | 46 | 5jmr | 5jmo | 71 | 2xxc | 2xxm |
| 22 | 6t2j | 6sc9 | 47 | 6o89 | 6o8d | 72 | 6jb9 | 1zvy |
| 23 | 6t2j | 6sc7 | 48 | 5vnv | 5vnw | 73 | 6jb9 | 6jb8 |
| 24 | 6t2j | 6sc8 | 49 | 4dka | 4dk6 | 74 | 5e03 | 5e5m |
| 25 | 6t2j | 6sc6 | 50 | 4dka | 4dk3 | 75 | 5ivo | 5ivn |
| | | | | | | 76 | 6apo | 6app |

**Table S1.** Pairs of PDB IDs for comparison of CDR3 compactness for bound and unbound nanobodies.

| | Molecular | | | | |
|---|---|---|---|---|---|
| 1 | Molecular Weight | 29 | Folded charge pH7 | 56 | Total atom-atom interactions |
| 2 | Sequence Length | 30 | Positive charge heterogeneity | 57 | Total carbon-pi interactions |
| 3 | Average Residue Weight | 31 | Negative charge heterogeneity | 58 | Total cation-pi interactions |
| | Amino acid composition | 32 | Isoelectric point (sequence) | 59 | Total donor-pi interactions |
| 4 | Aromatic content | 33 | Folded isoelectric point | 60 | Total sulfur-pi interactions |
| 5 | Tiny content | 34 | Unfolded isoelectric point | 61 | Total atom-plane interactions |
| 6 | Small content | 35 | Hydrophobicity | 62 | Total amide-amide interactions |
| 7 | Aliphatic content | 36 | Hydrophobic moment | 63 | Total amide-ring interactions |
| 8 | Nonpolar content | 37 | Solubility | 64 | Total plane-group interactions |
| 9 | Polar content | | Photochemical | 65 | Total planar interactions |
| 10 | Basic content | 38 | Molecular Extinction Coefficient | 66 | Average pairwise distance of interacting atoms (Å) |
| 11 | Acidic content | 39 | Molecular Extinction Coefficient (Cysteine bridges) | 67 | Total interactions |
| 12 | Total free cysteines | 40 | % Molecular Extinction Coefficient | | Secondary structure |
| 13 | Total cysteine bridges | 41 | % Molecular Extinction Coefficient (Cysteine bridges) | 68 | Phi ($\phi$) angle (degrees) |
| | Electrochemical | | Stability | 69 | Psi ($\psi$) angle (degrees) |
| 14 | Charge at pH1 | 42 | Instability Index | 70 | Total residues in isolated $\beta$-bridges |
| 15 | Charge at pH2 | 43 | Aliphatic Index | 71 | Total residues in $\beta$-turns |
| 16 | Charge at pH3 | | Interactions | 72 | Total residues in $\beta$-strands |
| 17 | Charge at pH4 | 44 | Total covalent interactions | 73 | Total residues in 3-10 helices |
| 18 | Charge at pH5 | 45 | Total van der Waals clashes | 74 | Total residues in $\alpha$-helices |
| 19 | Charge at pH6 | 46 | Total van der Waals interactions | 75 | Total residues in bends |
| 20 | Charge at pH7 | 47 | Total proximal interactions | 76 | Total residues in loops |
| 21 | Charge at pH8 | 48 | Total hydrogen bonds | | Thermodynamic |
| 22 | Charge at pH9 | 49 | Total weak hydrogen bonds | 77 | Free energy of folding |
| 23 | Charge at pH10 | 50 | Total ionic interactions | | Solvent accessibility |
| 24 | Charge at pH11 | 51 | Total aromatic interactions | 78 | Solvent accessible surface area (SASA) |
| 25 | Charge at pH12 | 52 | Total hydrophobic interactions | 79 | Side chain SASA |
| 26 | Charge at pH13 | 53 | Total carbonyl group interactions | | Druggability |
| 27 | Charge at pH14 | 54 | Total polar interactions | 80 | Structural Aggregation Propensity (SAP) |
| 28 | Unfolded charge pH7 | 55 | Total weak polar interactions | 81 | Developability Index (DI) |

**Table S2.** Additional *in silico* descriptors calculated for our nanobody datasets grouped by class according to Bashour *et al.* [27].

| Samples | CIC Retention Time (min) | HIC Retention Time (min) | BVP-ELISA OD450 (a.u.) | AC-SINS Δλmax (nm) | DLS PDI | Total CDR length | CDR3 length | CDR3 compactness | Patches CDR Surface Hydrophobicity | Patches CDR Positive Charge | Patches CDR Negative Charge |
|---|---|---|---|---|---|---|---|---|---|---|---|
| Brivekimig1_VHH1 | 9.25 | 13.35 | 1.95 | 26.45 | 0.41 | 33 | 16 | 1.35 | 81.36 | 0.26 | 0.22 |
| Brivekimig1_VHH2 | 9.24 | 13.37 | 0.23 | 27.18 | 0.36 | 33 | 16 | 1.38 | 94.24 | 0.06 | 0.17 |
| Brivekimig2_VHH1 | 9.38 | 16.60 | 3.16 | 7.25 | 0.09 | 24 | 8 | 0.83 | 106.99 | 0.00 | 0.00 |
| Caplacizumab_VHH1 | 9.33 | 13.34 | 0.22 | 26.61 | 0.62 | 37 | 21 | 1.53 | 92.06 | 0.57 | 1.22 |
| Enristomig_VHH1 | 9.23 | 15.23 | 0.11 | 4.38 | 0.26 | 20 | 5 | 0.56 | 79.36 | 0.05 | 0.00 |
| Enristomig_VHH2 | 9.39 | 11.80 | 1.02 | 26.58 | 0.97 | 30 | 15 | 1.04 | 106.50 | 0.06 | 0.00 |
| Envafolimab_VHH1 | 9.28 | 14.97 | 1.13 | 6.61 | 0.80 | 37 | 21 | 1.61 | 102.37 | 0.05 | 1.14 |
| Erfonrilimab_VHH2 | 9.48 | 20.87 | 3.31 | 5.03 | 0.22 | 37 | 21 | 1.57 | 102.99 | 0.00 | 0.00 |
| Gacovetug_VHH1 | 9.08 | 14.34 | 3.18 | 2.54 | 0.22 | 30 | 15 | 1.27 | 97.05 | 0.00 | 0.17 |
| Gefurulimab_VHH1 | 9.67 | 15.50 | 0.24 | 28.59 | 0.89 | 32 | 17 | 1.42 | 97.04 | 0.06 | 0.18 |
| Gefurulimab_VHH2 | 9.53 | 16.24 | 0.09 | 7.44 | 0.23 | 35 | 19 | 1.52 | 122.87 | 0.18 | 0.06 |
| Gocatamig2_VHH1 | 9.70 | 18.74 | 0.29 | 9.43 | 0.73 | 26 | 11 | 0.82 | 138.68 | 0.00 | 1.11 |
| Gontivimab_VHH1 | 9.37 | 19.24 | 1.93 | 6.15 | 0.17 | 35 | 19 | 1.36 | 99.40 | 0.04 | 0.11 |
| Gontivimab_VHH2 | 9.37 | 19.18 | 0.72 | 5.71 | 0.14 | 35 | 19 | 1.36 | 91.42 | 0.04 | 0.11 |
| Isecarosmab_VHH1 | 9.18 | 14.33 | 0.23 | 4.95 | 0.12 | 33 | 17 | 1.45 | 86.44 | 0.36 | 1.22 |
| Isecarosmab_VHH2 | 9.51 | 17.81 | 0.33 | 6.03 | 0.16 | 24 | 8 | 0.80 | 96.49 | 0.00 | 0.00 |
| Letolizumab_VHH1 | 9.49 | 16.97 | 1.39 | 9.48 | 0.87 | 27 | 11 | 1.00 | 95.13 | 0.27 | 0.28 |
| Lofacimig_VHH1 | 9.31 | 18.90 | 3.27 | 2.92 | 0.10 | 39 | 23 | 1.49 | 155.47 | 0.00 | 0.00 |
| Lofacimig_VHH2 | 8.89 | 19.84 | 2.30 | 1.79 | 0.31 | 35 | 20 | 1.32 | 119.68 | 0.00 | 1.53 |
| Lunsekimig1_VHH1 | 10.63 | 17.06 | 3.25 | 27.24 | 0.51 | 32 | 16 | 1.23 | 73.40 | 0.00 | 0.00 |
| Lunsekimig1_VHH2 | 9.96 | 16.88 | 3.23 | 15.21 | 0.10 | 35 | 19 | 1.39 | 116.89 | 0.00 | 0.00 |
| Lunsekimig2_VHH1 | 9.20 | 18.28 | 0.24 | 3.97 | 0.18 | 26 | 11 | 0.85 | 116.19 | 0.00 | 0.05 |
| Lunsekimig3_VHH1 | 8.89 | 13.38 | 0.17 | 2.04 | 0.14 | 36 | 18 | 1.48 | 97.22 | 0.10 | 1.88 |
| Ozekibart_VHH1 | 9.44 | 15.78 | 0.17 | 14.49 | 0.86 | 32 | 16 | 1.29 | 79.67 | 0.21 | 0.00 |
| Ozoralizumab_VHH1 | 9.69 | 14.04 | 0.27 | 29.78 | 0.08 | 24 | 8 | 0.98 | 83.65 | 0.10 | 0.00 |
| Ozoralizumab_VHH2 | 9.65 | 17.21 | 0.19 | 5.09 | 0.21 | 24 | 8 | 0.85 | 111.02 | 0.00 | 0.00 |
| Podentamig1_VHH1 | 9.31 | 15.12 | 3.28 | 4.21 | 0.53 | 28 | 14 | 0.92 | 86.31 | 0.02 | 0.00 |
| Podentamig1_VHH2 | 9.43 | 17.09 | 0.11 | 18.5 | 0.13 | 24 | 8 | 0.83 | 98.59 | 0.00 | 0.00 |
| Porustobart_VHH1 | 9.76 | 16.62 | 0.11 | 18.36 | 0.85 | 28 | 13 | 1.11 | 103.61 | 0.16 | 0.00 |
| Produvofusp_VHH1 | 9.07 | 17.10 | 1.23 | 3.61 | 0.12 | 38 | 22 | 1.57 | 90.60 | 0.00 | 0.29 |
| Rimteravimab_VHH1 | 9.06 | 16.9 | 0.15 | 3.99 | 0.14 | 34 | 18 | 1.43 | 89.51 | 0.00 | 0.65 |
| Sonelokimab1_VHH1 | 9.41 | 17.11 | 0.09 | 11.74 | 0.71 | 32 | 16 | 1.32 | 104.10 | 0.07 | 0.24 |
| Sonelokimab2_VHH1 | 9.55 | 16.38 | 0.83 | 3.23 | 0.05 | 31 | 17 | 1.35 | 97.97 | 0.23 | 1.45 |
| Tarperprumig_VHH1 | 9.96 | 16.72 | 0.22 | 30.35 | 0.94 | 32 | 17 | 1.42 | 99.13 | 0.06 | 0.14 |
| Tarperprumig_VHH2 | 9.22 | 14.86 | 0.17 | 3.68 | 0.24 | 28 | 13 | 0.89 | 99.60 | 0.30 | 0.14 |
| Vobarilizumab_VHH1 | 9.48 | 17.09 | 3.17 | 35.79 | 0.42 | 30 | 15 | 1.01 | 85.49 | 1.19 | 0.62 |

**Table S3.** Flag assignment and raw values for *in vitro* and TNP metrics for the 36 clinical-stage sequences. The raw data is available at *github.com/oxpig/TNP*.
